# Structure and ligand specificity of *Borrelia burgdorferi* BmpA

**DOI:** 10.64898/2026.01.30.702971

**Authors:** Qianqiao Liu, Victor A. Nuñez, Daniel Fernandez, Naima G. Sharaf

**Affiliations:** Department of Biology, Stanford University, Stanford, California, USA; Department of Structural Biology, Stanford University, Stanford, California, USA; Macromolecular Structure Group, Nucleus at Sarafan ChEM-H, Stanford University, Stanford, California, USA

**Keywords:** Borrelia burgdorferi, ABC transporters, substrate binding proteins, X-ray crystallography, bacterial lipoproteins

## Abstract

BmpA is a putative substrate-binding protein from *Borrelia burgdorferi*, the causative agent of Lyme disease, an organism with limited metabolic capacity that relies on salvage pathways rather than *de novo* nucleotide biosynthesis. Here, we determine the crystal structure of BmpA to a resolution of 2.6 Å, revealing a conserved substrate-binding protein fold with a deeply buried nucleoside-binding pocket. Using microscale thermophoresis, we show that BmpA binds thymidine with high affinity followed by cytidine and adenosine, whereas binding to ribose, guanosine, inosine, and uridine was not detected. Structure-guided mutagenesis further demonstrates that two conserved aromatic residues (Phe27 and Phe176) are essential for thymidine recognition, as alanine substitution at either position abolishes detectable binding. Additionally, a Foldseek-based structural homology search identified related proteins across diverse bacterial and archaeal species that share a conserved overall fold and binding-site architecture despite low sequence similarity, consistent with an evolutionarily conserved scaffold that can accommodate distinct nucleoside ligands. Together, our work illustrates how conserved binding protein architectures enable selective nucleoside acquisition and provides a foundation for understanding nutrient uptake strategies in organisms with reduced genomes.

## 1 Introduction

Lyme disease is caused by infection with the tick-borne pathogen *B. burgdorferi* sensu lato. After transmission, *B. burgdorferi* can replicate within the dermis (1), and early infection commonly presents with an expanding erythema migrans lesion, often described as a bull’s-eye rash (2). Without antibiotic treatment, the spirochete can disseminate to distal tissues, resulting in inflammatory manifestations including arthritis, carditis, and neurologic complications (3). Notably, the *B. burgdorferi* genome lacks many key protein-coding genes required for *de novo* synthesis of fatty acids, amino acids, nucleotides, and essential enzyme cofactors, reflecting a strong dependence on environment-derived nutrients (**Figure 1.A)** (4).

**Figure 1:**
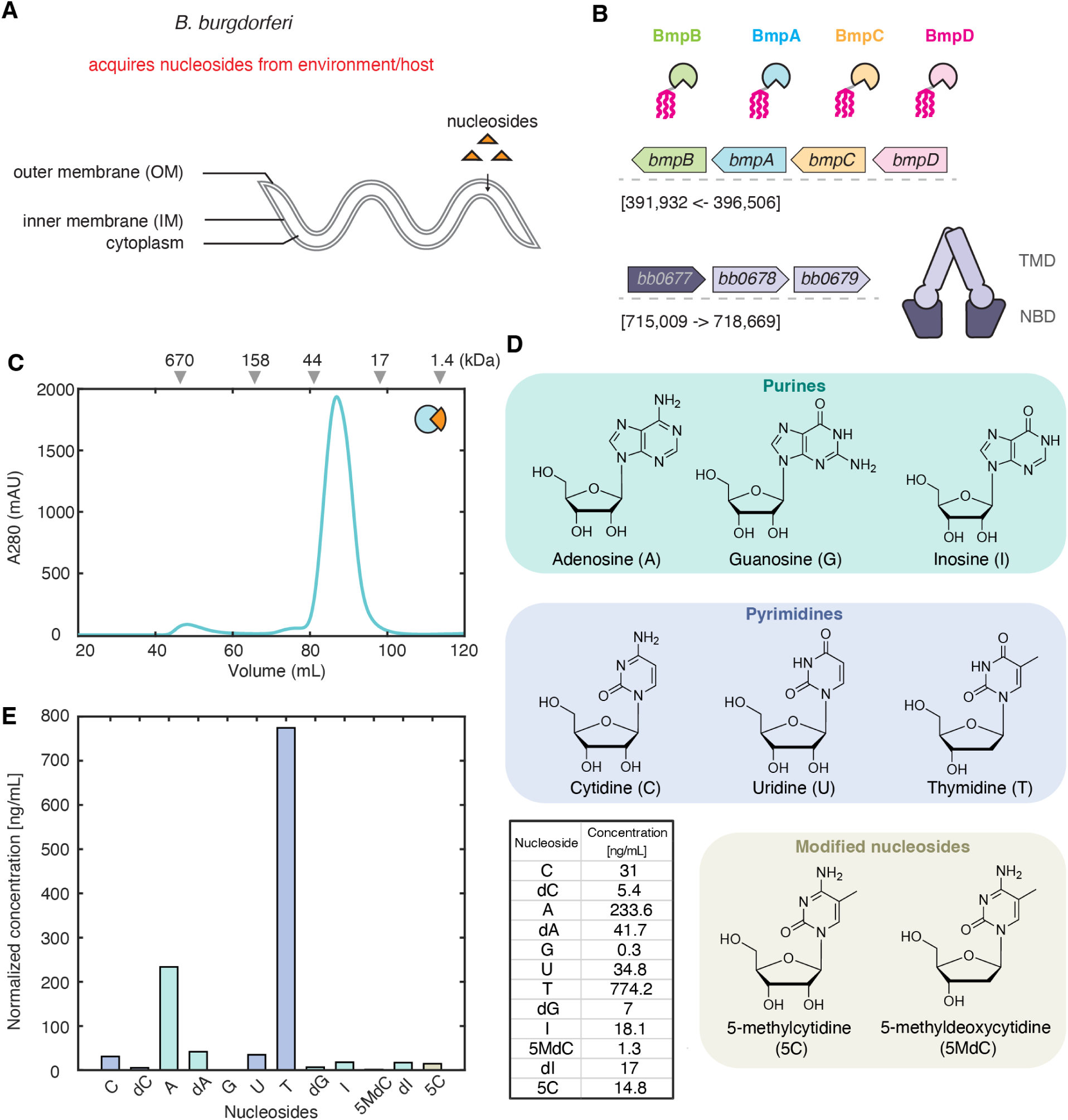
BmpA is a lipoprotein that co-purifies with endogenous nucleosides. **A.** Schematic of *B. burgdorferi* cell envelope architecture highlighting its reliance on host/environment-derived nucleosides. **B.** Genomic organization of the bmp locus and the putative ABC transporter operon encoding predicted nucleotide-binding domains (NBD) and transmembrane domains (TMD). Map coordinates are shown in brackets. **C.** SEC trace of purified BmpA. Inset: cartoon representation of BmpA with endogenous ligand. **D.** Detection of bound ligands by liquid chromatography–mass spectrometry. Molecular structures and their abbreviations of detected nucleosides are group by class. **E.** Relative abundance of nucleosides co-purifying with BmpA.

BmpA (basic membrane protein A), previously referred to as P39, is a lipoprotein that elicits a strong immune response (5). Antibodies to P39 have been detected as early as 7 days post-infection in animal models (6) and within the first month in human sera (7). Due to its high immunogenicity, it was found to be one of the most common specific markers for Lyme disease (8) and remains a diagnostically relevant antigen in current standard two-tier serological testing, where P39 is one of the proteins used to interpret immunoblots (9). The *bmp* locus in Borrelia burgdorferi comprises four paralogous genes: *bmpD*, *bmpC*, *bmpA*, and *bmpB* (**Figure 1.B)** (10). All four genes are transcribed *in vitro* and organized into two distinct transcriptional units. During growth *in vitro* Bmp genes are expressed in different amounts with bmpA transcribed at the higher level (65 fg of specific mRNA per 10 ng of total RNA), followed by bmpD and bmpB (each 20 fg) and bmpC (1.1 fg) (11).

To date, several studies have been conducted aimed at understanding the role of Bmp proteins in *B. burgdorferi* physiology. The importance of this family has been demonstrated using *bmpA/B* deletion strains which result in reduced joint colonization and attenuated arthritis. These phenotypes can be rescued by complementation (12). For BmpD, the structure and ligand-binding specificity has been characterized (13). Structural analysis of a BmpD crystal structure revealed that it adopts a canonical substrate-binding protein fold, typically seen for receptors of ATP-binding cassette (ABC) transporters. Ligand binding studies revealed that BmpD functions as a nucleoside-binding protein involved in the purine salvage pathway, with high specificity for guanosine and adenosine. This role in nutrient acquisition is important for *B. burgdorferi* since it lacks the enzymatic machinery for *de novo* biosynthesis of purines and pyrimidines (14).

*B. burgdorferi* is an c with host connective tissues, and *in vitro* it binds multiple extracellular matrix (ECM) components, including type I collagen, fibronectin and decorin (15–17). Consistent with this, ligand-affinity blot analyses of outer membrane–enriched fractions from the type strain B31 revealed several distinct laminin-binding proteins, including BmpA and its paralogs (18). This suggests an additional potential role for the Bmp family at the host–pathogen interface.

Despite BmpA’s prominence as a highly expressed member of the *bmp* locus and a diagnostically relevant antigen, the structure, ligand specificity, and molecular basis for substrate recognition remain poorly understood. Here, we determined the first crystal structure of BmpA at 2.6 Å resolution and show that it functions as a nucleoside-binding protein. Structural analysis complemented with mutational experiments reveal that ligand recognition is mediated by a conserved aromatic cradle that stabilizes the nucleobase and that surrounding residues play key roles in substrate specificity, providing a mechanistic framework for understanding how BmpA binds nucleosides in *B. burgdorferi*.

## 2 Results

### 2.1 Production of BmpA and characterization of endogenously bound ligands by mass spectrometry

Given that previous studies focused on BmpD, we sought to define the molecular basis for BmpA substrate recognition to enable a comparison within the Bmp family. To obtain sufficient amount of BmpA protein, we recombinantly expressed its soluble form in *Escherichia coli*. The signal peptide was removed, and an N-terminal hexahistidine tag was added for affinity purification purposes. BmpA was purified by nickel-affinity chromatography, followed by size-exclusion chromatography (SEC). Based on elution volumes relative to gel filtration standards, BmpA eluted at a molecular weight consistent with its monomeric forms on SEC and migrated at approximately 36 kDa on SDS-PAGE (**Figure 1.C**; **S1.A**).

Previous studies on BmpD identified adenosine as its endogenously bound ligand (13). To determine whether BmpA also has endogenously bound ligands, we performed liquid chromatography–mass spectrometry (LC-MS) analysis on the purified protein. As a control, we removed endogenously bound ligands from BmpA using an on-column unfolding-refolding protocol (**Figure S1.B-C**). We refer to this protein as uR-BmpA. As expected, only trace amounts of nucleosides were detected for uR-BmpA (**Figure S1.D**), suggesting that most endogenous ligands were removed by the unfolding/refolding protocol. In contrast, LC–MS analysis of BmpA detected multiple nucleosides with thymidine as the predominant bound species, followed by adenosine (**Figure 1.D-E**). These observations establish the endogenous ligand profile of BmpA when produced in *E. coli*.

### 2.2 Overall structure of BmpA

To understand the molecular basis of ligand binding, we solved the crystal structure of BmpA at a resolution of 2.6 Å (**Figure 2.A**). The structure was solved by molecular replacement using the crystal structure of BmpD (PDB ID:6SHU) as a search model. The asymmetric unit contained eight copies of the BmpA monomer. The structure shows that BmpA consists of two distinct domains linked by a flexible hinge region. The N-terminal domain consists of residues 8 to 115 and 243 to 269, and the C-terminal domain consists of residues 116 to 242 and 270 to 322. Each domain contains a central beta sheet composed of six *β*-strands, flanked by *α*-helices (**Figure S1.E**). In the N-terminal domain, the *β*-strands are arranged in a parallel orientation, whereas in the C-terminal domain, one strand (strand 12) is antiparallel to the others within the sheet. The crystal structure also revealed an endogenous ligand bound at the interface between the two domains. Based on the mass spectrometry data indicating that the BmpA sample was preferentially bound to thymidine, we modeled thymidine as the ligand (**Figure 2**, orange).

**Figure 2:**
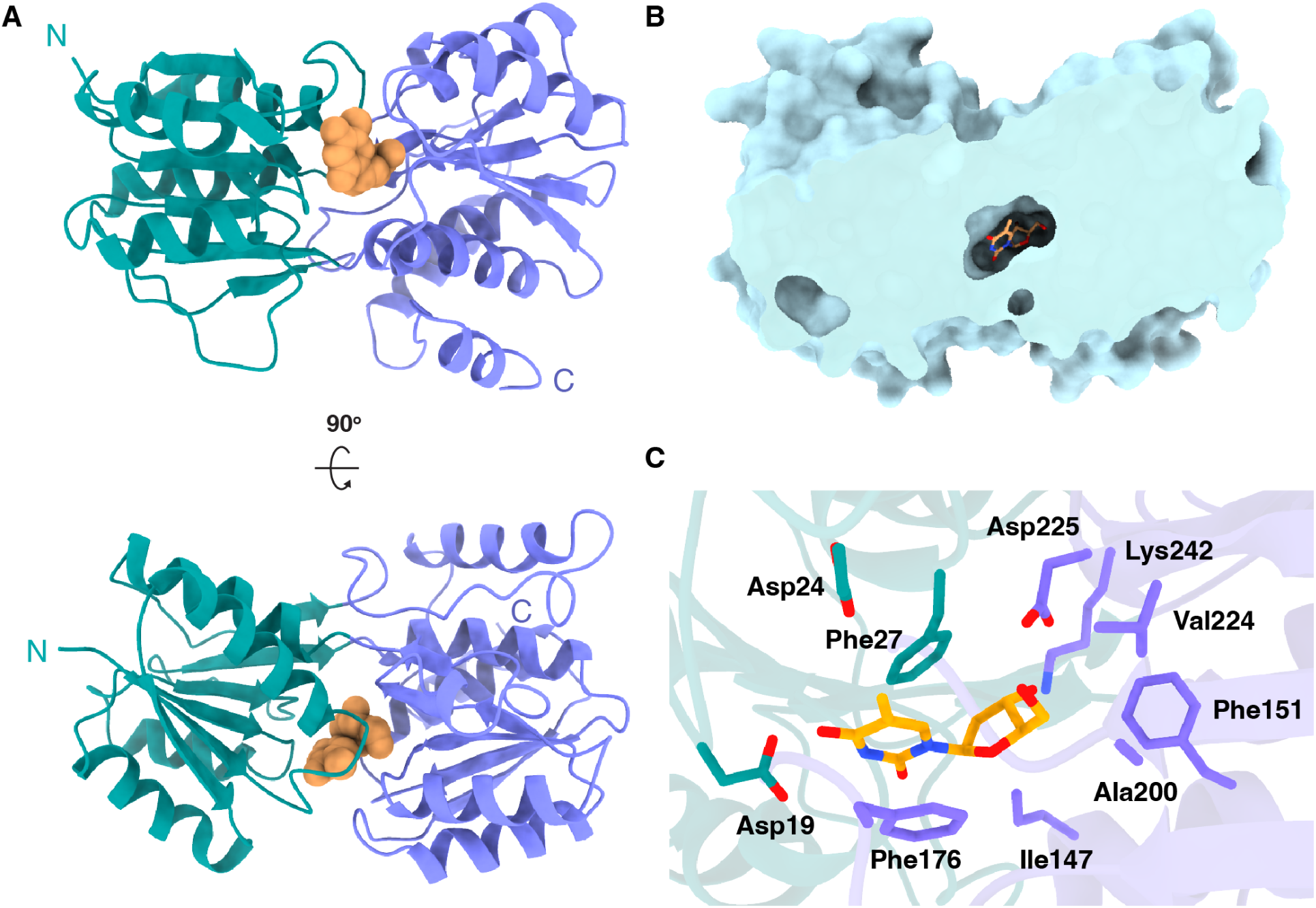
Structure of BmpA. **A.** Cartoon representation of BmpA shown in two orientation rotated by 90°. The N-terminal domain is colored teal, the C-terminal domain is purple. **B.** Surface representation of BmpA highlighting the ligand-binding cavity. **C.** Close-up view of the ligand-binding pocket in BmpA. BmpA backbone is in cartoon representation with key side chains highlighted as sticks. Thymidine is shown in orange for all panels.

### 2.3 Binding pocket analysis

Thymidine is bound in a discrete pocket lined by a mixed polar–hydrophobic environment (**Figure 2.B-C**). The polar face of the site is formed by Asp19, Asp24, and Asp225, with Lys242 positioned to provide complementary electrostatic interactions with thymidine’s polar groups. The other side of the cavity is dominated by hydrophobic residues. The planar pyrimidine ring packs against Phe27 and Phe176 (with additional hydrophobic enclosure from Phe151 and Ile147), stabilizing through van der Waals and *π*-associated contacts. Val224 and Ala200 further define the pocket boundary, shaping the cavity and constraining thymidine orientation. Together, these interactions delineate a binding site in which thymidine is anchored by a polar network and stabilized by an aromatic-rich hydrophobic cradle, providing a structural basis for nucleoside recognition by BmpA.

### 2.4 BmpA ligand binding analysis

To further characterize the binding affinity of BmpA for nucleosides, we performed MicroScale Thermophoresis (MST) using RED-labeled BmpA and a panel of ligands, including adenosine, cytidine, guanosine, thymidine, inosine, uridine, and ribose. Among these, BmpA exhibited the highest affinity for thymidine, followed by cytidine and adenosine, with dissociation constants (*K_d_*) of 5.32 *×* 10*^−^*^7^ M, 2.99 *×* 10*^−^*^6^ M, and 2.6 *×* 10*^−^*^5^ M, respectively (**Figure 3.A**). The remaining ligands, including uridine, which differs from thymidine by a single methyl and hydroxyl group, showed no measurable binding under the experimental conditions.

**TAble 1.** Data collection and refinement statistics.

| PDB ID | BmpA (9Q1U) |
| --- | --- |
| **Data collection <sup>a</sup>** |  |
| Wavelength (Å) | 0.97946 |
| Space group | P 2<sub>1</sub> 2<sub>1</sub> 2<sub>1</sub> |
| Cell dimensions |  |
| a, b, c (Å) | 106.27 133.17 191.39 |
| $\alpha$, $\beta$, $\gamma$ (°) | 90.00 90.00 90.00 |
| Unique reflections | 80623 (11985) |
| Multiplicity | 3.5 |
| Completeness (%) | 91.19 (90.35) |
| Mean I/sigma(I) | 4.5 |
| CC <sub>1/2</sub> % | 99.4 (15.6) |
| **Refinement** |  |
| Resolution (Å) | 50.60 - 2.58 |
| R-work/R-free | 0.254/0.306 |
| Number of non-hydrogen atoms | 14665 |
| macromolecules | 14490 |
| ligands | 113 |
| solvent | 62 |
| Protein residues | 1878 |
| RMS(bonds) | 0.14 |
| RMS(angles) | 0.33 |
| Ramachandran |  |
| favored (%) | 93.6 |
| allowed (%) | 5.5 |
| outliers (%) | 0.9 |
| Rotamer outliers (%) | 3.5 |
| Clashscore | 13 |
| Average B-factor, all atoms(Å<sup>2</sup>) | 86.0 |
| Numbers in parentheses correspond to the highest resolution shell |  |

**Figure 3:**
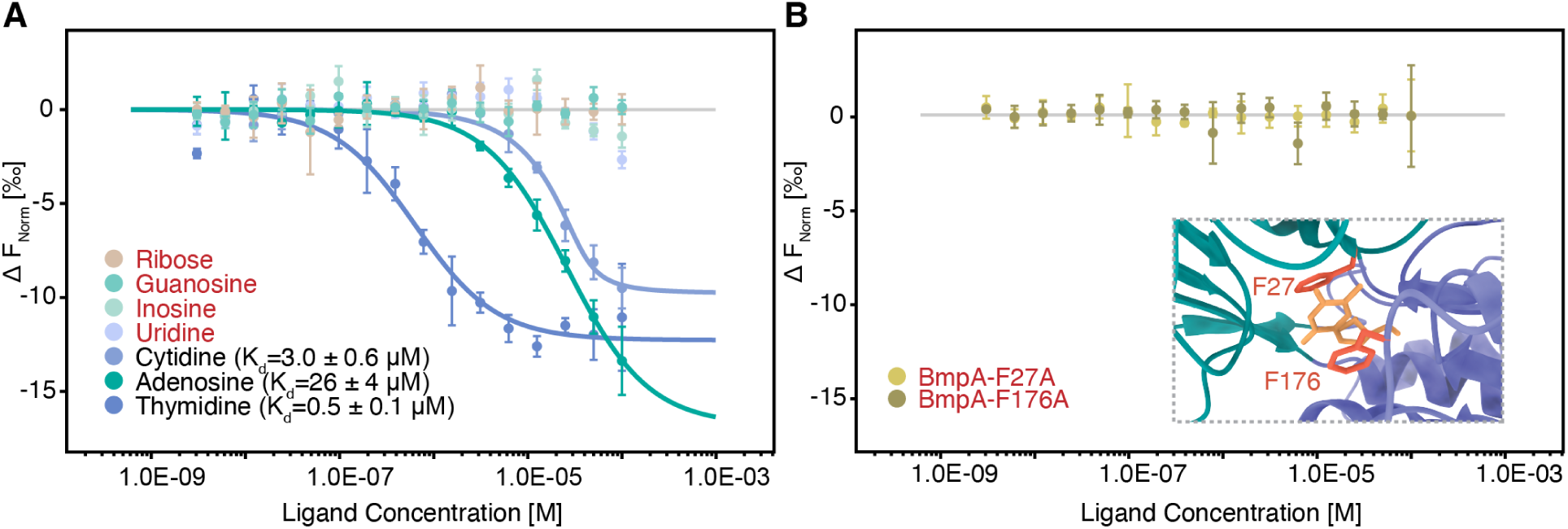
MST analysis of BmpA binding to ligands. **A.** Fitted curves of ligand binding to BmpA. The binding affinities of nucleosides to BmpA were determined by MST experiments. The concentration of NHS-RED-labeled BmpA was kept constant (10 nM), while the concentration of the non-labeled nucleosides ranged between 3.05 nM-100 *µ*M. **B.** MST binding analysis of thymidine with BmpA F27A and F176A mutants. The averages of three independent measurements are shown and error bars represent the standard deviation. Red labels indicate no detectable binding.

To investigate the molecular role of two phenylalanine residues (Phe27 and Phe176) in ligand binding, we generated alanine substitution mutants at these positions and assessed their ability to bind thymidine. Neither mutant exhibited detectable binding to thymidine, the highest-affinity ligand of wild-type BmpA (**Figure 3**.B). This complete loss of binding highlights the critical role of Phe27 and Phe176 in ligand recognition, underscoring their functional importance in the BmpA binding interface. Together, these findings demonstrate that BmpA exhibits specific and high-affinity binding to thymidine and that Phe27 and Phe176 are critical for this interaction. The results provide molecular insight into the ligand recognition mechanism of BmpA and suggest that these phenylalanine residues play key roles in the binding interface.

### 2.5 Comparison to Borrelia Bmp family and other substrate binding proteins

The genes *BmpA*, *BmpB*, *BmpC* and *BmpD* are located on the main chromosome of *Borrelia burgdorferi*, adjacent to each other. Protein sequence alignment reveals that BmpA shares less than 50% sequence identity with other members of the Bmp protein family, and BmpC is the most different among all (**Figure S2**). Despite their differences in sequence, these substrate binding proteins share a high structural similarity. The structural alignment of the crystal structures of BmpA and BmpD resulted in an RMSD of 0.9 Å. The two phenylalanines that stack the ligand are conserved in BmpA, BmpB, and BmpD. In BmpC, the first phenylalanie is replaced by tyrosine and the second one is replaced by leucine.

In BmpD, the adenosine ligand is stabilized by a network of hydrogen bonds and hydrophobic interactions involving residues such as Asp19, Phe27, Asn28, Asp225, and Phe176, which coordinate the nucleoside within a deep cavity (13). Similarly, the BmpA binding site maintains a conserved aromatic and polar environment. Although the overall fold remains highly similar, subtle differences in the arrangement and identity of ligand-coordinating residues, such as the substitution of specific aspartate and phenylalanine residues, may contribute to the distinct substrate preferences observed between BmpA and BmpD. These variations highlight the evolutionary adaptability of the nucleoside-binding pocket within this protein family and may help explain the differential binding affinities for nucleosides.

To compare BmpA to other structurally similar proteins, we performed a structural homology search using Foldseek (19). This analysis identified homologous proteins from diverse bacterial and archaeal species (13, 20–23), with varying degrees of sequence identity to BmpA (**Figure 4**). The overall fold and ligand-binding architecture are broadly conserved, despite low sequence similarity among some family members, underscoring both the evolutionary conservation and adaptability of this protein family in recognizing different nucleoside substrates. Across these structures, nucleoside ligands are accommodated deep within the binding pocket formed by conserved residues (Phe27, Phe151, Phe176, Val224, Asp225, and Lys242; numbering according to BmpA) (**Figure 5**). Notably, the structures contained distinct ligands, including *Steptococcus pneumoniae* PnrA bound to uridine, suggesting that differences within the binding pocket, rather than the overall fold, govern ligand recognition and specificity.

**Figure 4:**
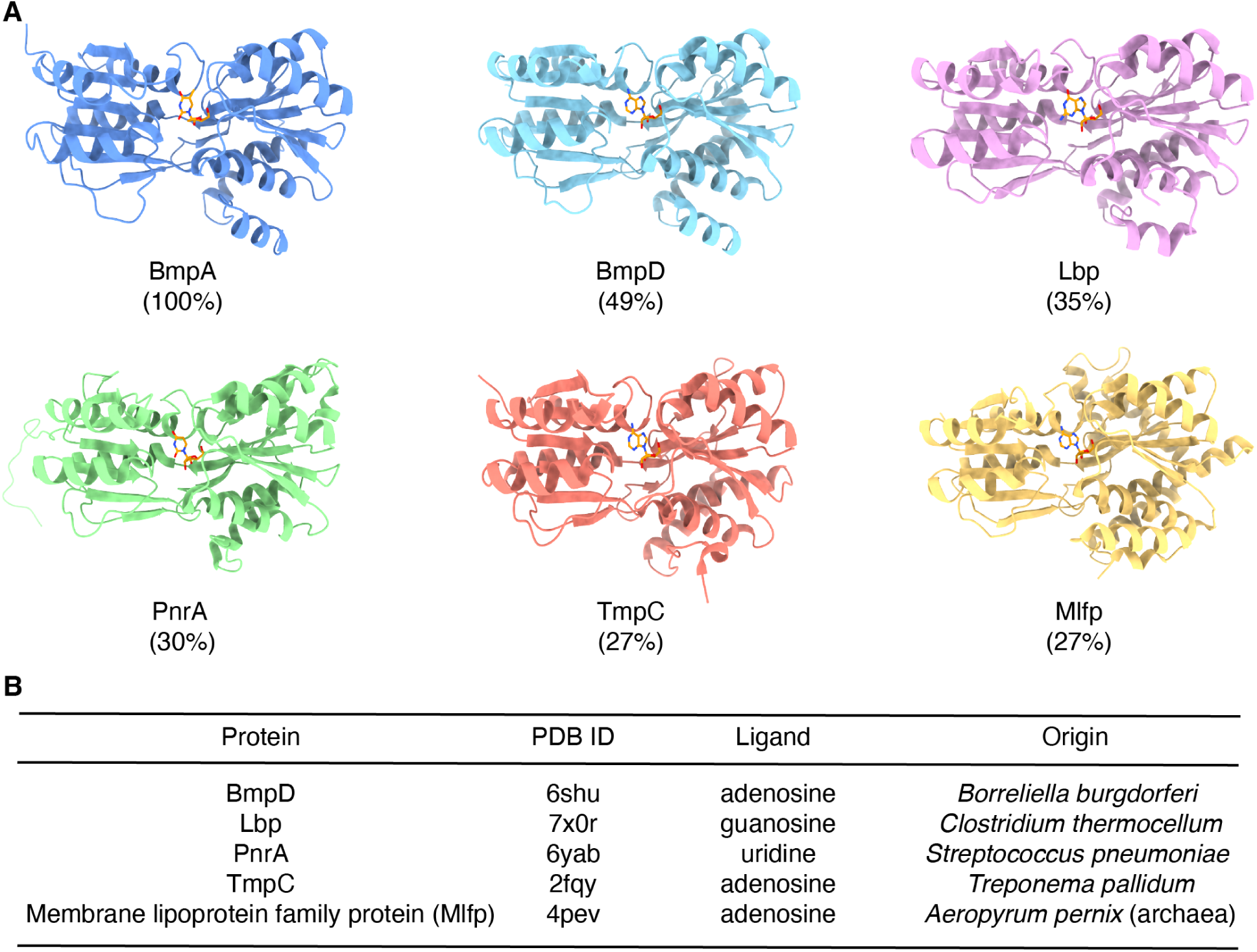
Comparison of BmpA to other substrate-binding proteins identified using Foldseek. **A.** Structures and percent sequence identities relative to BmpA. Each protein is shown in a distinct color: BmpA (blue), BmpD (cyan), Lbp (purple), PnrA (green), TmpC (salmon), and membrane lipoprotein family protein (yellow). Bound ligands are depicted in stick representation. **B.** Table summarizing each protein, PDB ID, ligand, and species of origin.

**Figure 5:**
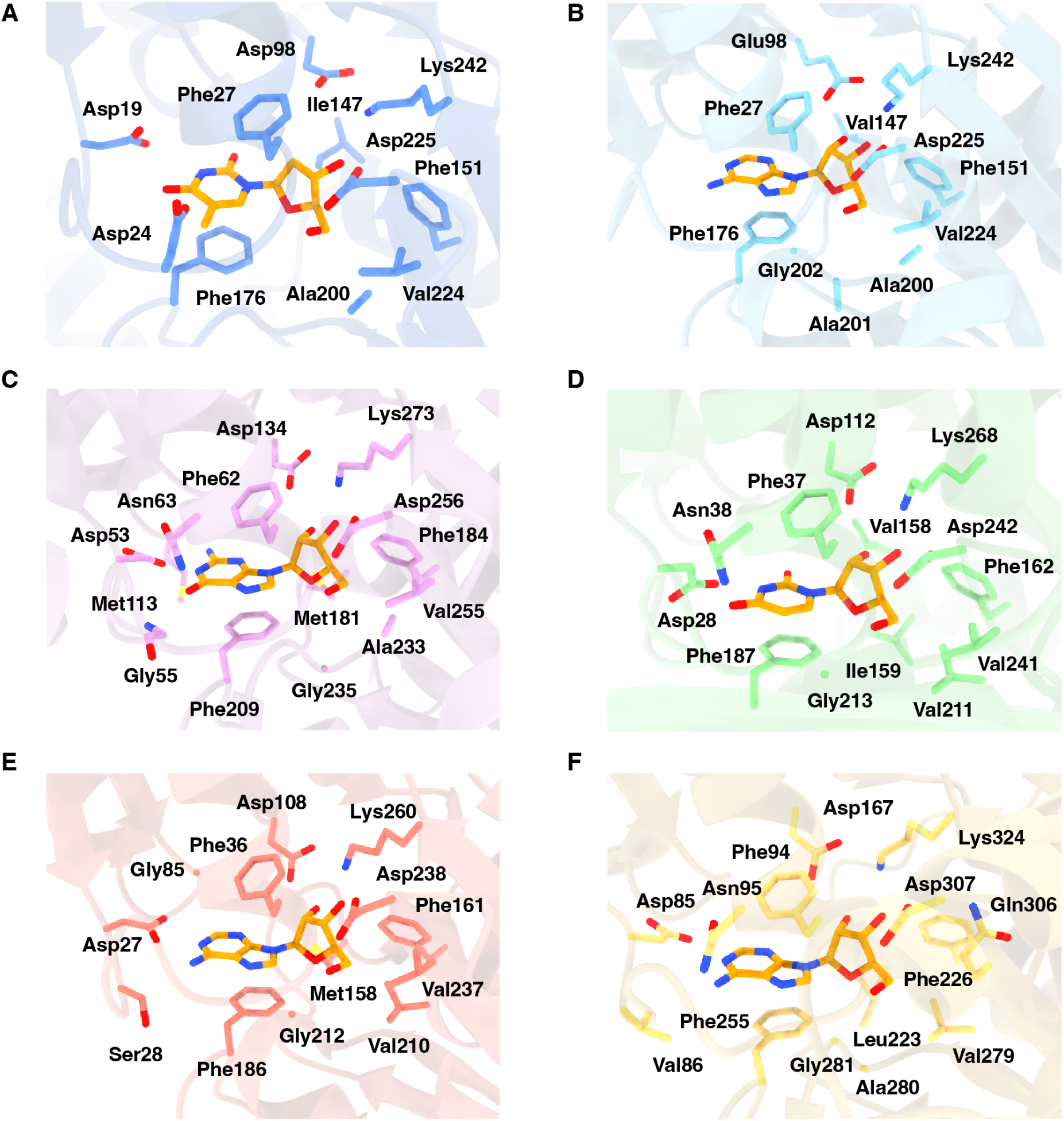
Ligand-binding sites for the BmpA structural homologs. Bound nucleosides are shown in stick representation, and binidng pocket residues are shown as sticks and labeled. Protein backbones are shown as cartoons (A-F: BmpA, blue; BmpD, cyan; Lbp, purple; PnrA, green; TmpC, salmon; Mlfp, yellow).

## 3 Discussion

Nutrient acquisition is essential for bacterial growth and survival. One notable feature of *B. burgdorferi* is its reduced genome, which only contains 1,291 protein-encoding genes, compared to 4,401 in the model organism *E. coli* K12 (24). Key protein-encoding genes with an apparent absence in the *B. burgdorferi* genome include those involved in the synthesis of fatty acids, amino acids, nucleotides, and enzyme cofactors (4). Without these proteins, *B. burgdorferi* is unable to synthesize many essential molecules *de novo*, a limitation highlighted by its need for rich media when grown under laboratory conditions (25). Therefore, to survive, *B. burgdorferi* must rely on its transport proteins to acquire nutrients from its environment. To gain structural insight into *B. burgdorferi* transport proteins we chose to characterize the structure and ligand specificity of BmpA, a putative substrate binding protein whose paralog (BmpD) has been characterized as a purine binding protein. Given the limited protein repertoire of *B. burgdorferi*, the presence of multiple Bmp paralogs is unexpected and suggests selective pressure for the production of multiple similar proteins.

To gain insights into the structure and ligand specificity of BmpA, we recombinantly expressed and purified the protein. LC-MS analysis provided an initial indication that thymidine was the predominant bound ligand, suggesting a preference for this nucleoside. This result contrasts with previous mass spectrometry analysis of BmpD, which showed preferential binding to adenosine (13). To further define BmpA ligand specificity, we performed MST binding assays, which showed that BmpA binds nucleosides with a clear preference for pyrimidines. Specifically, among the ligands tested, thymidine bound with the highest affinity, followed by cytidine and then adenosine, whereas other tested nucleosides showed no measurable binding under the assay conditions. Notably, ribose alone showed no detectable interaction, consistent with the requirement for a nucleobase and mirroring observations reported for BmpD (13).

To define the structural basis for this selectivity, we determined the crystal structure of BmpA in complex with thymidine. Here we report the first crystal structure of BmpA determined at 2.6 Å resolution. The BmpA crystal structure reveals a typical two lobe substrate binding protein-like fold with a deep ligand-binding pocket. Analysis of the thymidine-bound structure reveals a binding pocket that integrates a polar anchoring region with an aromatic-rich hydrophobic surface that packs against the nucleobase. Two phenylalanines, Phe27 and Phe176, sit at key positions within the pocket and are essential for ligand binding, as alanine substitution at either position abolished measurable thymidine binding. These results show that these residues are essential for complex formation with thymidine, likely by stabilizing the planar pyrimidine ring through van der Waals and *π*-associated interactions and by enforcing a productive nucleoside pose that allows polar contacts to the sugar/base heteroatoms. The lack of ribose binding further supports this model, as sugar alone cannot engage the aromatic platform that appears to be essential for nucleoside capture. Together, these data support a recognition mode in which these residues play key roles in the selection of nucleosides over sugars.

While our structural and mutational analyses provide a mechanistic explanation for nucleobase selectivity in BmpA, they also offer a framework for understanding how closely related SBP homologs discriminate among similar nucleosides. Comparison to other BmpA-like SBPs indicates that these proteins share a conserved overall fold and a broadly conserved ligand-binding architecture, but differ at a small number of positions within the binding pocket—differences that can be sufficient to shift ligand preference. An example is TmpC (PnrA; Tp0319) from *Treponema pallidum*, which displays a ligand preference opposite to BmpA: TmpC binds uridine (*K_d_ ≈* 13 *µ*M) but not thymidine (22). Despite this functional divergence, superposition of the TmpC and BmpA crystal structures reveals a highly conserved binding-site scaffold, including the aromatic “sandwich” that stabilizes the nucleobase (Phe27 and Phe176 in BmpA) and conserved polar residues positioned to engage the ribose (Asp98, Lys242, and Asp225 in BmpA). The most prominent differences are localized near the N-terminus (e.g., Asp27 and Ser28 in TmpC) and are positioned to influence local hydrogen-bonding patterns and loop positioning at the entrance of the pocket. Additional insight comes from PnrA in *Streptococcus pneumoniae*, which binds five nucleosides (adenosine, guanosine, cytidine, uridine, and thymidine) (21). Comparison of its nucleoside-bound structures indicates that specificity is tuned by the conformational plasticity of a short loop spanning residues 27–36 (corresponds to 26–36 in TmpC), which can rearrange to accommodate the distinct hydrogen-bonding and steric features of purine versus pyrimidine bases. Taken together, these comparisons support a model in which a conserved phenylalanine-based aromatic cradle provides a shared platform for nucleobase recognition, whereas substrate selectivity is encoded by subtle, local variations, particularly in flexible loops and nearby polar side chains, superimposed on an otherwise conserved SBP scaffold. This framework suggests that closely related Bmp proteins could evolve distinct nucleoside specificities with minimal changes to global structure.

A central challenge for *B. burgdorferi* is maintaining nutrient acquisition across the diverse microenvironments encountered during its enzootic cycle, including transitions between the tick vector and animal host. The physiological importance of the Bmp proteins is underscored by prior work demonstrating that deletion of *bmpA/bmpB* genes results in impaired growth in mouse joint tissue (12). In mice, *bmpA* and *bmpB* are dramatically and selectively upregulated in joints at days 12 and 15 post-infection compared with skin, heart, and bladder, whereas *bmpC* and *bmpD* show no tissue-specific differences and remain unchanged in joints over the same time frame. In this context, our work provides a mechanistic explanation as to why *B. burgdorferi*, despite its reduced genome, encodes multiple Bmp proteins: if the availability of nucleoside varies across vector and host microenvironments, divergent ligand specificity among Bmp proteins could ensure robust nutrient uptake under fluctuating metabolic conditions encountered during infection and transmission, with BmpA specialized for pyrimidines and BmpD for purines. Future direct measurements of nucleoside concentrations in relevant host and vector microenvironments, paired with *in vivo* expression profiling of the *bmp* genes, will be crucial to test this model experimentally.

In addition to its role in nutrient acquisition, BmpA may also function as a surface-associated factor. As a diderm bacterium, *B. burgdorferi* can localize lipoproteins to either the inner or outer membrane, where they may either face the periplasm or be surface-exposed. Localization studies of the Bmp proteins have yielded seemingly contradictory results. On one hand, BmpA, BmpB, BmpC, and BmpD have all been shown to bind laminin, a component of the extracellular matrix (18), and antibodies have detected BmpA and BmpB on the bacterial surface (12). These studies suggest that the Bmp proteins are localized on the surface. On the other hand, studies using recombinantly overexpressed proteins suggest that BmpB, BmpC, and BmpD are associated with the inner membrane (26). Interestingly, this study did not report BmpA localization. This omission is particularly surprising since BmpA, like all substrate binding protein involved in nutrient acquisition, must be present in the inner membrane to interact with its cognate ABC transporter. The data presented here demonstrate that BmpA binds nucleosides, strongly suggesting that it plays a role in nutrient acquisition. However, key gaps remain in firmly establishing this model. To date, there are no reports either confirming direct interaction between BmpA and its putative transporter or its localization at the inner membrane (**Figure 6**). It therefore remains possible that BmpA binds nucleosides at the outer membrane without playing a direct role in nutrient uptake at the inner membrane.

**Figure 6:**
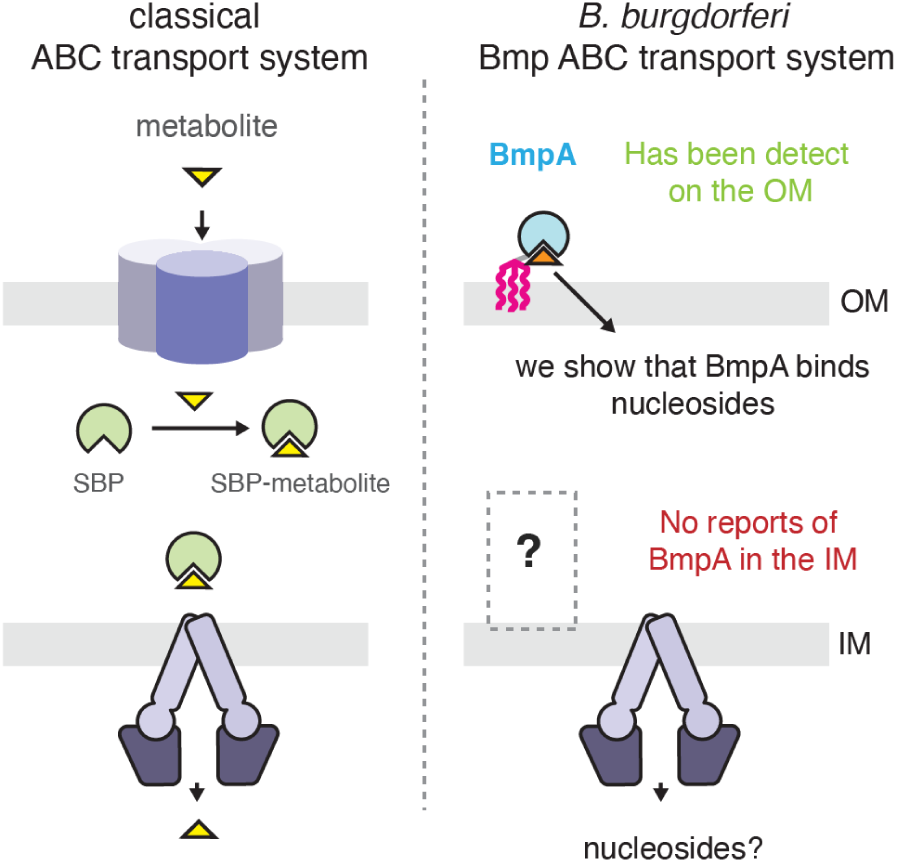
Comparison of *B. burgdorferi* Bmp-associated ABC transport system and a classical ABC transporter system. Classical ABC transport systems use a periplasmic SBP to capture a metabolite and deliver it to an inner-membrane transporter for transport into the cell (Left). A working model for the Bmp system in *B. burgdorferi* (Right).

Alternatively, BmpA may exhibit dual localization and function, acting both as surface-associated adhesin and as a substrate binding protein involved in nutrient acquisition at the inner membrane. Such “moonlighting” behavior is not unprecedented in *B. burgdorferi* : the surface-exposed 5’-methylthioadenosine*/S* -adenosylhomocysteine nucleosidase (MTAN; Bgp) exhibits both metabolic activity in nutrient salvage and a role in glycosaminoglycan binding implicated in host-cell adherence, and MTAN inhibitors can impair spirochete growth (27). Moreover, similar examples exist in other pathogens, including *Neisseria meningitidis*, where the methionine substrate binding protein NmMetQ is a lipoprotein that binds multiple methionine analogs, stimulates the ATPase activity of its cognate importer, and exhibits dual localization consistent with a “moonlighting” role at the cell surface (28). Dissecting localization-dependent functions and establishing experimental evidence directly linking BmpA to its cognate ABC transporter will be an important direction for future work.

In summary, we determined the first crystal structure of BmpA, and our analysis of this structure provided mechanistic insights into ligand recognition and specificity. Specifically, the thymidine-bound pocket reveals an aromatic cradle in which Phe27 and Phe176 are essential determinants of thymidine recognition, while comparative analyses reveal that changes within the ligand binding pocket may tune ligand specificity. Because BmpA and the previously characterized BmpD have different substrate specificities, we suggest that these protein perform complementary roles in nucleoside uptake. This division of substrate specificity may expand nucleoside salvage capacity across environments and provides a mechanistic framework for understanding why *B. burgdorferi*, despite its limited protein repertoire, encodes both of these proteins. Together, this study provides insight into nutrient acquisition and establish a mechanistic framework for future antimicorbial strategies targeting nucleoside import in *B. burgdorferi*.

## 4 Experimental procedures

### 4.1 Cloning, expression, and purification of BmpA proteins

The amino acid sequence for *Borrelia burgdorferi* BmpA was obtained from the B31 strain (UniProt ID: Q45010). The coding region for BmpA was amplified and ligated into the pET21b(+) vector (ampicillin resistance) between the NdeI and XhoI restriction sites, placing expression under control of the T7 promoter (GenScript Biotech). The signal peptide (residues 1–18) was removed and an N-terminal hexahistidine tag was added for purification. The recombinant plasmid was transformed into *E. coli* BL21(DE3) gold cells (Agilent Technologies). Cultures were grown in ZYM-5052 autoinduction medium (containing 100 mg/L ampicillin) at 30°C for 30 hours. Cells were harvested by centrifugation at 4,875 RCF for 15 minutes at 4°C (JLA 8.1 rotor; 5000 rpm), and the cell pellets were flash frozen in liquid nitrogen and stored at -80°C until further use.

For purification, approximately 10 g of cell paste was thawed and resuspended in 100 mL of buffer containing 25 mM Tris–HCl pH 7.5, 100 mM NaCl, 40 mg lysozyme, 4 mg DNase, and one cOmplete protease inhibitor cocktail tablet (Sigma-Aldrich). Cells were disrupted using a Microfluidizer (Microfluidics), and lysates were clarified by centrifugation at 7,823 RCF (Ti45 rotor; 10,000 rpm) for 30 min at 4°C. Proteins were purified using a 5 mL HisTrap High Performance column (Cytiva), pre-equilibrated with 25 mM Tris–HCl pH 7.5 and 100 mM NaCl (buffer A). BmpA was eluted with a gradient of up to 300 mM imidazole in buffer A. The collected protein fractions were further purified by size-exclusion chromatography with a Hiload 16/600 Superdex 200 pg column (Cytiva), equilibrated in buffer A. The on-column unfolding and refolding procedure was done as previously described (29). Peak fractions were pooled, snap frozen in liquid nitrogen, and stored at -80°C. Protein purity and identity were confirmed by SDS-PAGE.

### 4.2 Crystallization, structure determination and analysis

Recombinant BmpA were brought to a final concentration of 30 mg/ml and prepared for crystallization experiments. Crystallization trials were conducted at 16°C by the sitting-drop vapor diffusion technique, utilizing conditions from the MBClass II Suite (Molecular Dimensions). The crystals formed in drops containing 0.1 M sodium chloride, 0.1 M sodium acetate, 0.1 M lithium sulfate, and 12% (w/v) polyethylene glycol 4000. For cryoprotection, individual crystals were flash-soaked for a few seconds in a mixture of reservoir solution supplemented with 20% (v/v) glycerol prior to being vitrified in liquid nitrogen.

X-ray diffraction data were collected at SSRL beamline 12-1 at the Stanford Synchrotron Radiation Lightsource (SSRL) at the SLAC National Accelerator Laboratory, and processed using the XDS software package (30). The crystals of BmpA belonged to the P2_1_ P2_1_ P2_1_ space group. The structure was determined by molecular replacement with PHASER (31), using the structure of BmpD as the search template (PDB ID: 6SHU). Iterative model refinement was performed with Phenix-1.20.1 and Coot 0.9.8.7 (32, 33). The final structure factors and coordinates were deposited in the Protein Data Bank under accession code 9Q1U.

Protein secondary structure diagrams were generated using PDBsum and annotated with ESPript3.0 (34, 35). Amino acid sequence alignments were conducted through UniProt’s alignment tool and presented using ESPript3.0. Gene locations of Bmp proteins were obtained from the BioCyc database (36). Analysis and rendering of BmpA electron density map and structure were done in ChimeraX-1.9 (37). To define the interaction interface, the Contacts tool in ChimeraX was employed with its default parameters (VDW surface overlap cutoff at -0.4 Å). Structural homologues of BmpA (9Q1U) were identified by searching the PDB100 database using Foldseek (19). All figure assembly and annotation were completed in Adobe Illustrator 2025.

### 4.3 LC-MS analysis

HRMS data at both MS1 and MS2 levels were acquired on a Thermo Fisher Scientific Q Exactive HF-X orbitrap (RRID:SCR 018703) monitor and confirm structures of nucleosides. Once analyte structures were confirmed, a sensitive and robust LC-MS/MS method using a Waters Xevo TQ-XS triple quadrupole mass spectrometer (RRID:SCR 018510) was applied using a one-point calibration strategy to estimate concentrations of analytes using a standard mixture of known concentration. Liquid chromatography using Waters Acquity LC system was performed on a Phenomenex Synergi 4um Fusion-RP 80 Å column with an acetonitrile/water/0.1% formic acid mobile phase and gradient elution with a total run time of 15 minutes. Detection and quantification were carried out by positive heated electrospray ionization (HESI). To account for differences in ionization efficiency, peak areas for each component were normalized to those of corresponding 50 ng/mL standard solutions.

### 4.4 MicroScale Thermophoresis

Binding interactions between BmpA and select ligands were assessed using microscale thermophoresis. Purified BmpA was fluorescently labeled with NHS-RED dye following the manufacturer’s protocol (NanoTemper Technologies). The labeled BmpA was diluted to a final concentration of 10 nM in PBST buffer (phosphate-buffered saline, 0.05% Tween-20, pH 7.4). A series of ligand dilutions was prepared to achieve a concentration range from 100 *µ*M to 3.05 nM. Each MST reaction consisted of mixing equal volumes of NHS-RED-labeled BmpA and nucleoside solution, followed by a 10-minute incubation at room temperature to ensure binding equilibrium. Prepared samples were loaded into standard MST premium capillaries (NanoTemper Technologies). Measurements were taken using a Monolith LabelFree/pico RED instrument set to 100% excitation power and medium MST power. All experiments were performed in triplicate. Data analysis was performed using MO.Affinity Analysis software (NanoTemper Technologies). Binding affinities (Kd) were determined by fitting the normalized fluorescence responses against the logarithm of ligand concentration using a 1:1 binding model.

## 5 Acknowledgments

This work was funded by the Howard Hughes Medical Institute Emerging Pathogens Initiative program. We gratefully acknowledge Ludmila Alexandrova and the Stanford University Mass Spectrometry Laboratory for assistance with LC–MS method development, data acquisition, and analysis. We thank Silvia Russi at the Stanford Synchrotron Radiation Lightsource (SSRL) Beamline 12-1. Use of the Stanford Synchrotron Radiation Lightsource, SLAC National Accelerator Laboratory, is supported by the U.S. Department of Energy, Office of Science, Office of Basic Energy Sciences under Contract No. DE-AC02-76SF00515. The SSRL Structural Molecular Biology Program is supported by the DOE Office of Biological and Environmental Research, and by the National Institutes of Health, National Institute of General Medical Sciences (P30GM133894). The contents of this publication are solely the responsibility of the authors and do not necessarily represent the official views of NIGMS or NIH. Microscale thermophoresis analyses were performed at the Macromolecular Structure Group at the Nucleus, Stanford University (RRID: SCR 023233). This work utilized the NanoTemper Monolith Labelfree/pico RED system that was purchased with funding from Stanford c-SHARP Program (RRID:SCR 022986).

## Supporting Information Available

**Figure S1:**
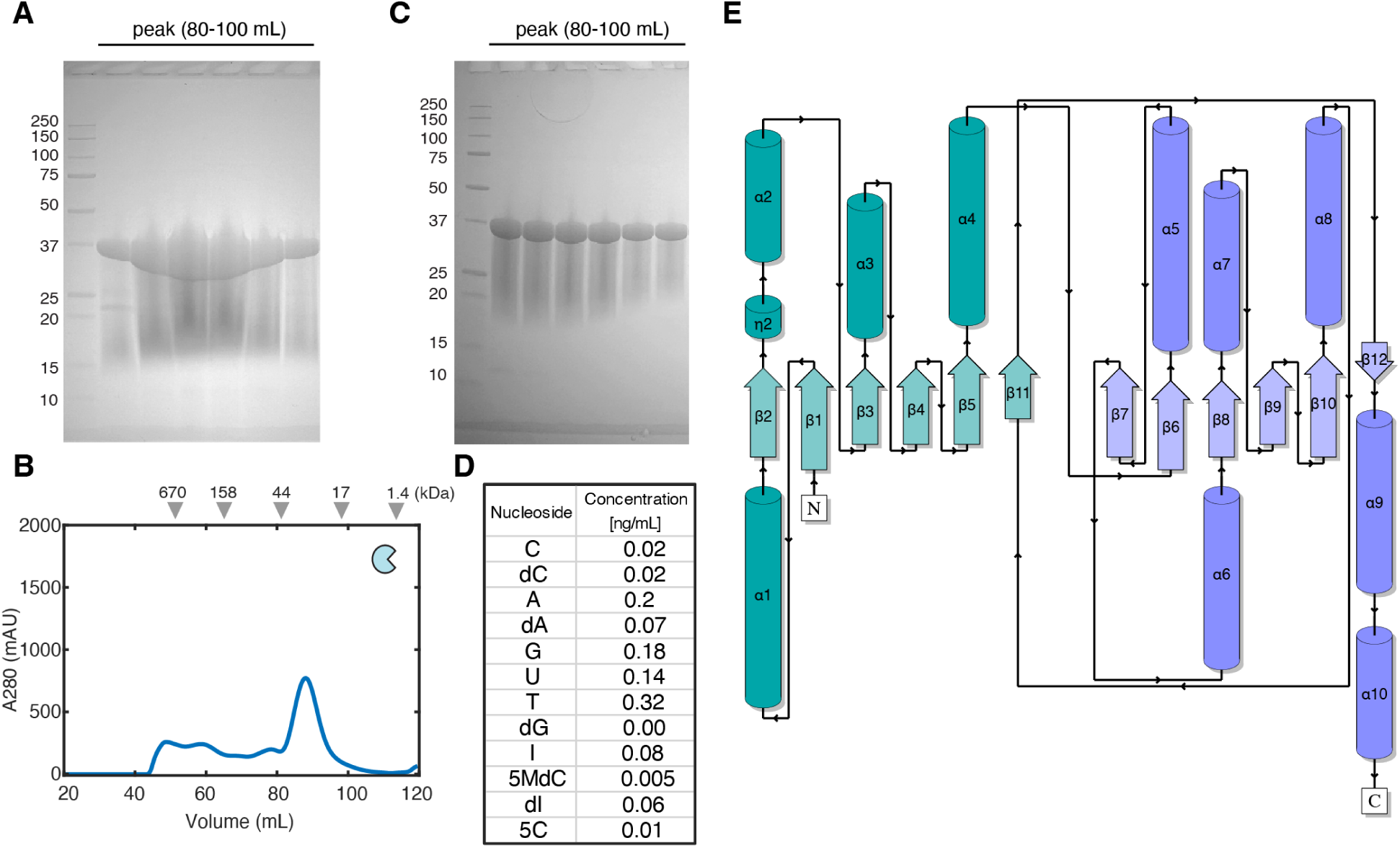
**A.** SDS-PAGE of purified BmpA protein. **B.** SEC trace of unfolding/refolding treated BmpA protein. **C.** SDS-PAGE of unfolding/refolding treated BmpA protein. **D.** LC-MS analysis of unfolding/refolding treated BmpA. **E.** Schematic of BmpA secondary structure. The N-terminal domain is colored teal and the C-terminal domain is purple.

**Figure S2:**
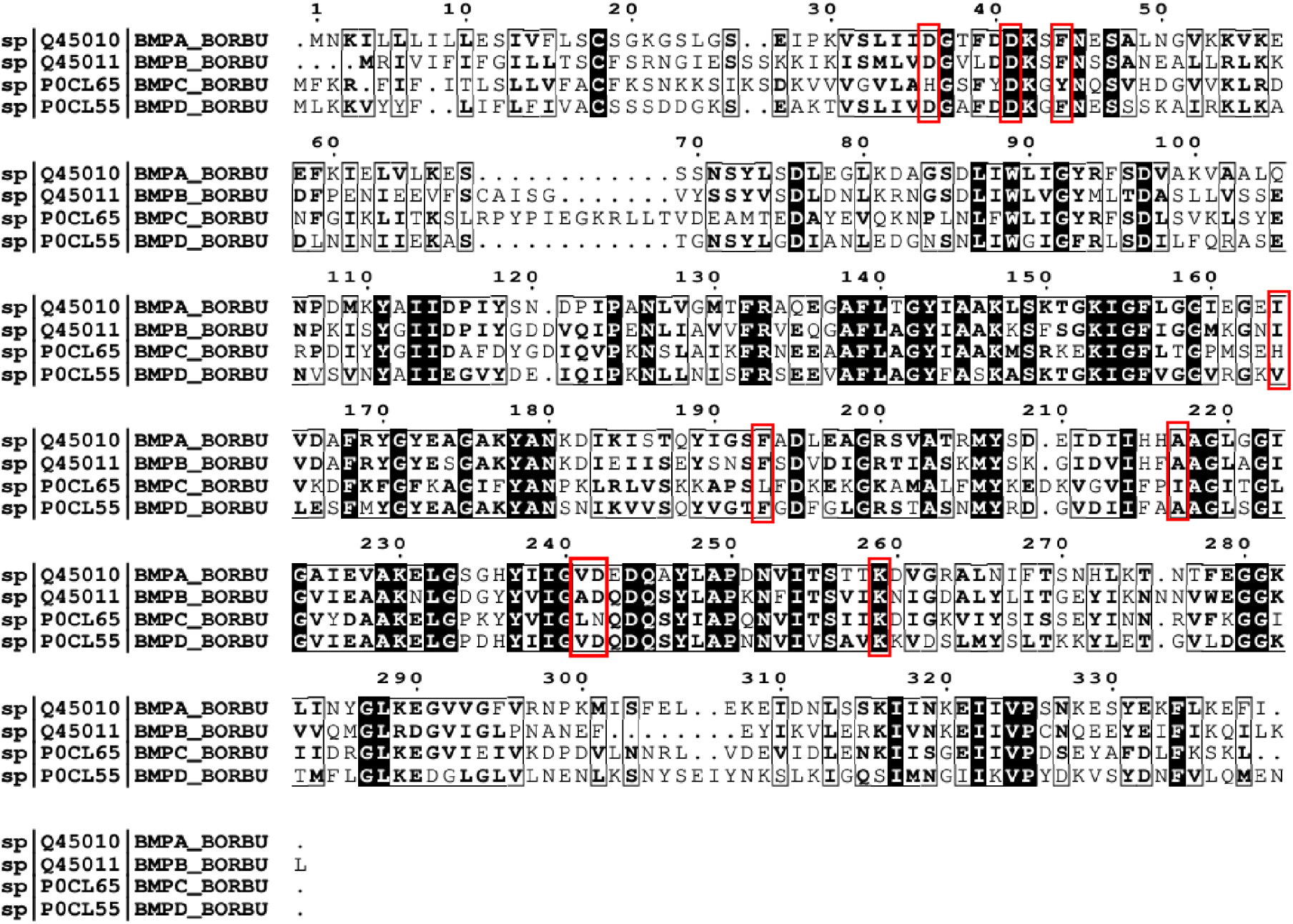
Sequence alignment of BmpA with BmpB, BmpC and BmpD. White characters in a black box indicate strict identity, bolded characters indicate similarity within a group, a black frame indicates similarity across groups, and residues involved in ligand contact are boxed in red. Residue numbering is based on BmpA.

